# Novel combination of irreversible electroporation and allogenic chimeric antigen receptor (CAR) T-cell therapy synergizes therapeutic outcomes in a preclinical human pancreatic cancer mouse model

**DOI:** 10.1101/2025.08.10.669537

**Authors:** Edward Jacobs, Julio Arroyo, Sam Salemizadeh Parizi, Wei Guo, Yong Lu, Rafael Davalos

**Author notes:** Corresponding Author, 313 Ferst Drive NE, Atlanta, GA, USA, 30318. These authors contributed equally (Edward Jacobs, Julio Arroyo).

## Abstract

Irreversible electroporation (IRE) is a non-thermal ablation modality used clinically for treating unresectable tumors while preserving vital structures through controlled application of pulsed electric fields. Previous data suggest that patient outcomes are enhanced with the induction of an anti-tumor immune response, but current research focuses on using immune checkpoint inhibitors, which function through conventional immune pathways that may be downregulated by cancer or dysregulated by chemo-induced lymphodepletion. Chimeric Antigen Receptor (CAR) T-cells overcome this limitation, as they are engineered with synthetic receptors that redirect lymphocytes to recognize and target cells expressing tumor-specific structures. CARs are engineered to have an increased binding affinity compared to in-situ T-cell binding, amplify internal stimulation cascades, and release pro-inflammatory cytokines that can modulate the endogenous immune system. However, there are still major limitations for adoptive cell therapies in solid tumors, including life-threatening on-target off-tumor cytotoxicity, antigen escape, and failure to infiltrate and persist in solid tumors. Given the substantial evidence that IRE overcomes many of the challenges associated with immune infiltration and persistence in solid tumors, there is a strong premise for using targeted cell therapies following IRE, which would then target residual cancer that could repopulate the lesion. Here, we present the first proof-of-concept combination of IRE with an adoptive cell therapy. We validated that the cell membrane CAR target is not affected in electroporated cells that survive IRE, allowing for subsequent binding and elimination of residual tumor. The research demonstrates the feasibility and synergy of a novel combination of two clinically used techniques.

## Introduction

Pancreatic cancer is the 7^th^ leading cancer-related death worldwide, with a 5-year survival rate of ∼13%^1^. Despite decades of work improving surgical procedures, chemoradiation, and early diagnostic techniques, pancreatic cancer is still an insidious prognosis due to its surreptitious and aggressive growth. Consequently, most pancreatic cancer patients are diagnosed at locally advanced or metastatic stages, with 80 – 90% having pancreatic ductal adenocarcinoma (PDAC)^2^.

PDAC is characterized by substantial vascular and ductal involvement that precludes surgical resection in >80% of patients^3^. Further, even for amenable patients who undergo surgical resection, >50% experience local tumor recurrence within a year^4^.

Irreversible electroporation (IRE) is a non-thermal focal pulsed field ablation technique used clinically for the treatment of unresectable and aggressive tumors in the prostate^5–7^, kidney^8–10^, liver^11^, and pancreas^12,13^. IRE employs high-voltage (1000–3500 V) microseconds-long (70–100 µs) monophasic pulses, applied through thin surgical probes placed within or adjacent to the tumor^14,15^. The electric field generated within the tissue from the applied voltage permeabilizes the cellular membrane through the formation of nano-scale pores (electroporation)^16–18^. If persistent, cells die due to the inability to maintain homeostasis^19^. Due to the non-thermal mechanisms, IRE can ablate large volumes of tissue (>50cm^3^) without significantly heating the surrounding tissue or structures^20,21^, allowing it to be delivered near the bowel^22^, ducts^23^, mature blood vessels^24,25^, and nerves^26,27^. Thus, IRE is one of the few treatments available when current surgical resection and thermal ablation methods are contraindicated. Consequently, IRE nearly triples the median survival of patients diagnosed with stage III locally advanced pancreatic cancer (LAPC) from 6-13 months to 17-44 months^28–37^.

The poor tumor location for pancreatic cancer is paralleled by extensive genetic heterogeneity, an impenetrable stroma, and an immunosuppressive “cold” tumor microenvironment (TME), which limits the localization and persistence of targeted therapies^38^. Multiple cell populations contribute to the immunosuppressive TME, including differentiated cancer cells, cancer stem cells, tumor-associated fibroblasts, and immunosuppressive immune cells (e.g., tumor-associated macrophages, myeloid-derived suppressor cells, and regulatory T-cells)^39^. Pancreatic cancer stem cells account for 0.5-1% of the tumor cell population in PDAC and exhibit an increased resistance to chemoradiation and targeted therapies, allowing escape from therapeutic intervention^40,41^. Further, the epigenetic and cellular composition of tumors can vary between patients, between different tumors within a patient, and even at different locations within the same tumors^42^, making it challenging to provide single-target therapeutics. IRE acts indiscriminately on proliferating and non-proliferating cells within the lethal electric field^43^, producing sub-millimeter demarcations between ablated and non-ablated tissue. Therefore, recalcitrant (e.g., cancer stem cells^47,48^) and immunosuppressive immune cells are removed in addition to bulk tumor cytoreduction^44–46^. IRE also alters the physical properties of the TME by reducing extracellular matrix density and rigidity^52,53^, as well as increasing tumor-associated blood vessel permeability^47–51^. Together, this reverses the stroma-induced immunosuppression and tumor-associated hypoxia, thereby potentiating immune cell infiltration and immunotherapy delivery to the cancer^47^.

Destruction of the cellular membrane through IRE also releases damage-associated molecular patterns, which initiate the innate inflammatory response, and tumor-associated antigens^52^. Unlike thermal ablation modalities, the antigens released are not denatured due to the lack of sufficient thermal heat, leading to antigen-mediated T-cell activation through phagocytosis and presentation by antigen-presenting cells^53^. While overall survival and progressive-free survival for LAPC have been significantly extended with IRE, curative outcomes are still limited by local and distant recurrence^54–56^. Though treatment success is not predicated by the induction of a pro-inflammatory response, immune activation correlates with survival outcomes^29^. Both the innate and adaptive immune systems are demonstrated to activate and localize within the TME following IRE^46^.

However, systemic anti-tumor responses following IRE are not consistent between patients, which may lead to eventual tumor recurrence. Many patients receive chemotherapy before and/or after IRE treatment^33,35,57^, which can induce lymphodepletion and weaken the adaptive T-cell response^58^. In an attempt to more consistently generate a persistent peripheral anti-tumor immune response, IRE has predominantly been combined with immunotherapies (*i.e.*, anti-CTLA- 4, -PD-L1, and -PD-1)^45,47,52,54^. In a mouse model, primary tumor ablation followed by anti-CTLA4 and anti-PD1 immune checkpoint inhibitors significantly increased tissue-resident and circulating memory cytotoxic T-cells targeting the cancer-specific antigen, SPAS-1^52^. Subsequently, this work indicated that a tumor vaccine effect was achieved by the tissue-resident and circulating memory cytotoxic CD8^+^ T-cells, limiting the reintroduction of new cancer. A recent direct comparison of IRE with cryoablation and thermal ablation further demonstrated that anti-PD1 synergizes best with IRE, leading to longer tumor-free survival, increased infiltration of CD8^+^ T- cells, and protection against tumor reintroduction^59^. He *et al*. presented promising clinical results when combining IRE with anti-PD1 in LAPC, achieving an overall survival of 44.3 months compared to the 23.4 months for IRE alone^54^.

To overcome aspects of the lymphodepletion and lack of endogenous T-cell priming, immune cells can also be removed from the patient before chemoradiation, primed *ex vivo*, then infused back^60,61^. Delivery of natural killer (NK) cells and allogenic γδ T-cells has been briefly demonstrated concomitant with IRE in humans^60,62,63^, but NK cells do not generate a sustained or specific anti-tumor immune response^64^. Further, while γδ T-cells are less restricted by specific tumor antigens, potentially enabling broader cancer recognition, major histocompatibility complex I (MHC I) is significantly downregulated in many cancer types, precluding the binding of primed T-cells, with PDAC patients specifically exhibiting suppression rates of 40-100%^65,66^. Without receptors for immune cell recognition, micrometastases are hidden from the heightened immune response and eventually repopulate local and distant sites. Chimeric Antigen Receptor (CAR) T-cells overcome this limitation, as they are engineered with synthetic receptors that redirect T lymphocytes to recognize and target cells expressing tumor-specific surface structures, proteins, and glycans, while the T-cell Receptor (TCR)-MHC complex targets processed intracellular proteomes^67,68^. CARs have been engineered to have an increase in binding affinity over *in-situ* TCR-MHC binding, and generation II – IV CARs have internal signal cascade amplification, which can lead to higher functionality over conventional TCR pathway activation^69,70^. Further, activated CAR T-cells release pro-inflammatory cytokines that modulate the endogenous immune system^67,71,72^, sometimes with side effects when excessive^73^. Though CAR-T therapy has achieved unprecedented success in hematopoietic cancers^74^, there are still major limitations that must be addressed, including life-threatening on-target off-tumor cytotoxicity, antigen escape, and failure to infiltrate and persist in solid tumors^73^. There are also well-defined pathways that inhibit CAR T-cell immunity and recruitment within the TME, including anti-inflammatory cytokines and chemokines released by immunosuppressive immune cells, physical barriers created by the tumor-recruited fibroblasts, and immune inhibitors^73^.

Given the substantial evidence that IRE overcomes many of the challenges associated with endogenous T-cell infiltration and persistence in solid tumors that are shared with CAR-T therapy, there is a strong premise for using targeted adoptive cell therapies following IRE, which we hypothesize would then target residual cancer that could repopulate the lesion. Here, we demonstrate a proof-of-concept combination of irreversible electroporation with human CAR T-cell therapy. We demonstrate that the cell membrane CAR target is not affected in electroporated cells that survive IRE, allowing for subsequent binding and elimination of residual tumor. The research presented demonstrates scientific and clinical feasibility for the novel combination of two actively used clinical techniques.

## Results

### Local tumor recurrence is indicated following irreversible electroporation of pancreatic cancer in an immunodeficient mouse model

Both IRE and CAR T-cell therapy are immunogenic, with the immune system influencing the response to primary and metastatic disease. Tumor response following IRE is enhanced in immunocompetent mice compared to immunodeficient nude mice^75^, and recent work indicates that IRE produces an abscopal effect in an immunocompetent contralateral Pan02 pancreatic cancer C57Bl/6 mouse model, with the size of the secondary tumors remaining stable following treatment of the primary tumor^76^. However, many patients receive chemoradiation that weakens the immune response; therefore, we investigated tumor control following IRE in immunodeficient Pan02-bearing NOD/SCID/IL2gc-KO (NSG) mice. NSG mice lack functional T, B, and NK cells, allowing for the assessment of IRE without adaptive immune-mediated cytotoxicity of residual disease. Though cell death is indicated <24 hours after IRE, the necrotic portions of the tumor need to be cleared by the innate immune system. Together with potential scabbing, the presence of small residual tumors cannot be measured with calipers, so we transfected the Pan02s to express Firefly Luciferase for *in vivo* imaging system (IVIS) use to assess the treatment response *via* tumor luminescence^77^.

We first utilized our Pan02-laden collagen hydrogels to quantify the lethal threshold for irreversible electroporation (Figure 1A, 525 ± 77 [442 – 613] V/cm) and then simulated the expected ablative tumor coverage for subcutaneous tumors with randomized conductivities and sizes (Figure 1B), as previously detailed^78,79^. We calculated the percent tumor coverage for applied voltage-to-distance ratios ranging from 1000 to 3000 V/cm and determined that a voltage-to-distance ratio of 2500 V/cm should fully ablate every tumor (Figure 1C). Informed by the computational model, each tumor was then treated with adjustments to the electrode spacing and voltage made based on the measured tumor size at the time of treatment. A significant drop in luminescence was observed on IVIS imaging on days 3 and 7 (Figure 1D), indicating complete coverage of the tumor by the ablative field. All tumors were also visibly flattened and scabbed by day 3 post-treatment (Figure 1E), confirming tumor ablation and justifying the need for imaging to determine tumor response following pulsed electric field treatments. Despite tumor clearance indicated by imaging on days 3 and 7 post-treatment, scabbing was mostly healed, and tumor regrowth was evident in all mice and 66% (4/6) of tumors on day 17. The total measured photon flux within the tumor region of interest (Figure 1F), as determined from the IVIS images, and the tumor volume over time confirmed partial tumor regrowth in several mice following initial regression (Figure 1G).

**Fig 1.**
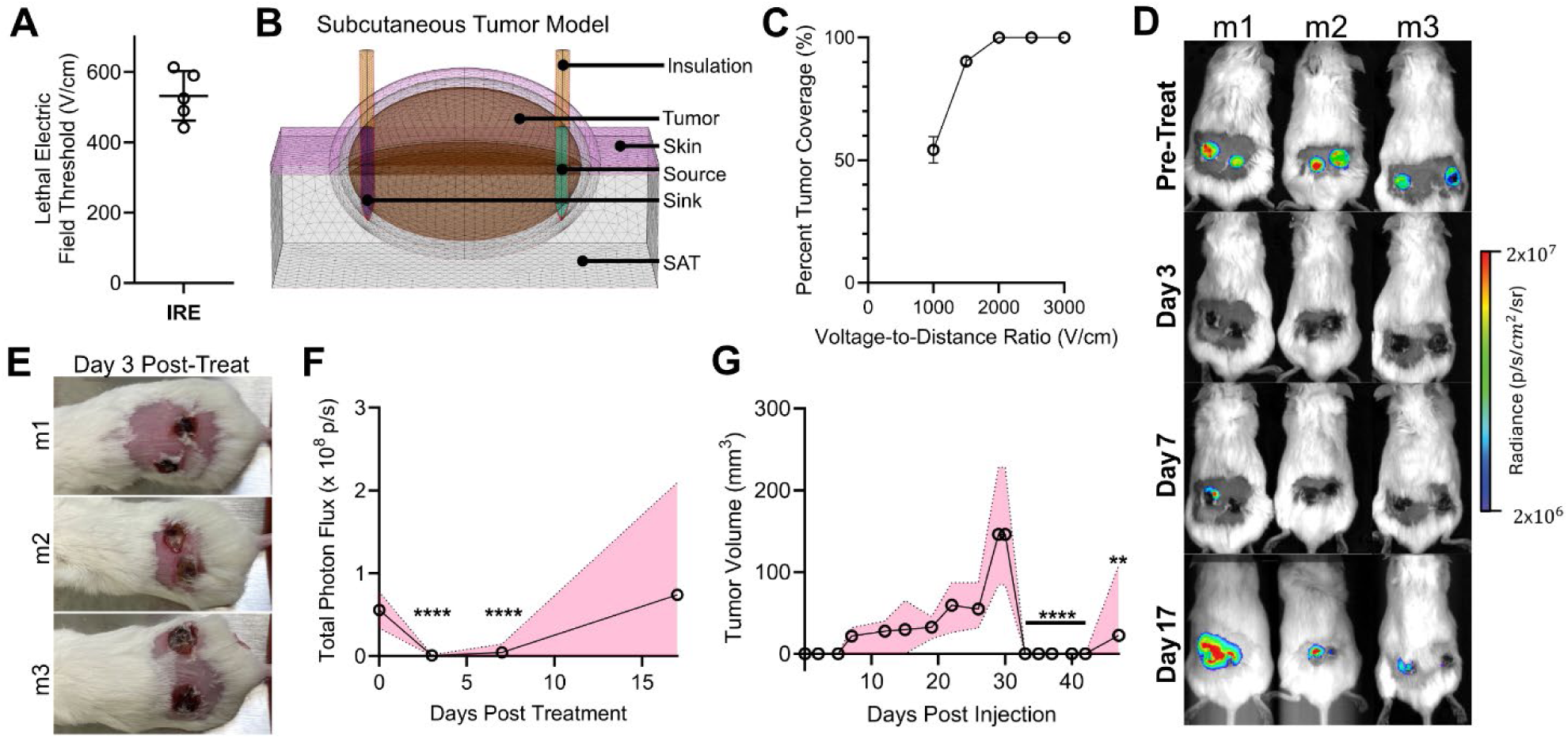
A) Calculated IRE lethal electric field thresholds for Pan02 mouse pancreatic cancer; mean ± SD; n =5. **B)** COMSOL Multiphysics finite element model (FEM) representing subcutaneous mouse tumors with 5-mm spacing shown. **C)** Simulated percent tumor coverage with a lethal electric field, using the subcutaneous FEM with randomized tissue conductivities and tumor sizes. **D)** *In vivo* imaging system (IVIS) of FLuc-eGFP^+^ Pan02s in immunodeficient NSG mice (m) follows complete ablation and subsequent microtumor recurrence after IRE treatment at 2500 V/cm; n=6. **E)** Tumor scabbing and flattening at day 3 post-treatment. **F)** Total photon flux within the tumor region of interest from the IVIS images; mean and range presented; Multiple two-tailed T-tests between the data compared to the initial day zero total photon flux; n=6. **G)** Measured tumor size over time from tumor inoculation; mean and range presented; Multiple two-tailed T-tests between the data compared to pre-treatment tumor size on day 30; n=6. ** *p <* 0.01, **** *p <* 0.0001.

Further, for all recurrent tumors, the growths appear to originate in a small region on the edge of the tumor. These findings suggest that while IRE induces acute cytoreduction in pancreatic tumors, residual viable cells can persist and repopulate the lesion in immunodeficient mice. These results underscore the critical role of immune surveillance in achieving durable treatment responses and motivate the development of adjunct immunotherapeutic strategies, such as CAR T-cell therapy, to eliminate post-IRE residual disease.

### CAR target binding is not affected in viable cells following irreversible electroporation

Due to the thinness of the cell membrane, the induction of a transmembrane potential during IRE pulse delivery can generate electric fields within the membrane on the order of ∼200 MV/m, large enough to potentially destabilize or denature membrane proteins^16,80,81^. Further, the organization and function of membrane proteins are also dependent on lipid interactions, and electroporation physically alters the cell membrane composition and structure^82–84^, possibly disrupting the recognizable binding motif within the lipoprotein complex. Therefore, to interrogate whether electroporation precludes the binding of CAR T-cells due to dysregulating membrane proteins, we evaluated mesothelin binding *via* flow cytometry following treatment. Mesothelin is a membrane protein that is commonly overexpressed in human pancreatic cancers and is consequently a frequent CAR target^85,86^ . We utilized the AsPC-1 human pancreatic cancer cell line, which is indicated to overexpress mesothelin and has been used for human CAR T-cell experiments^87,88^. IRE may disrupt cellular homeostasis, but residual cellular function can allow for the metabolism of viability-associated proteins; therefore, the cells were also stained to identify viable cells, with the gating set based on the control, non-treated AsPC-1 population (Figure 2B), to distinguish between high-viability and low-viability populations and between high-mesothelin and low-mesothelin expression. Compared to Jurkat human leukemia cells, which are negative for mesothelin (Figure 2C)^77^, the wild-type AsPC-1s were validated for high mesothelin expression (Figure 2D).

**Fig 2.**
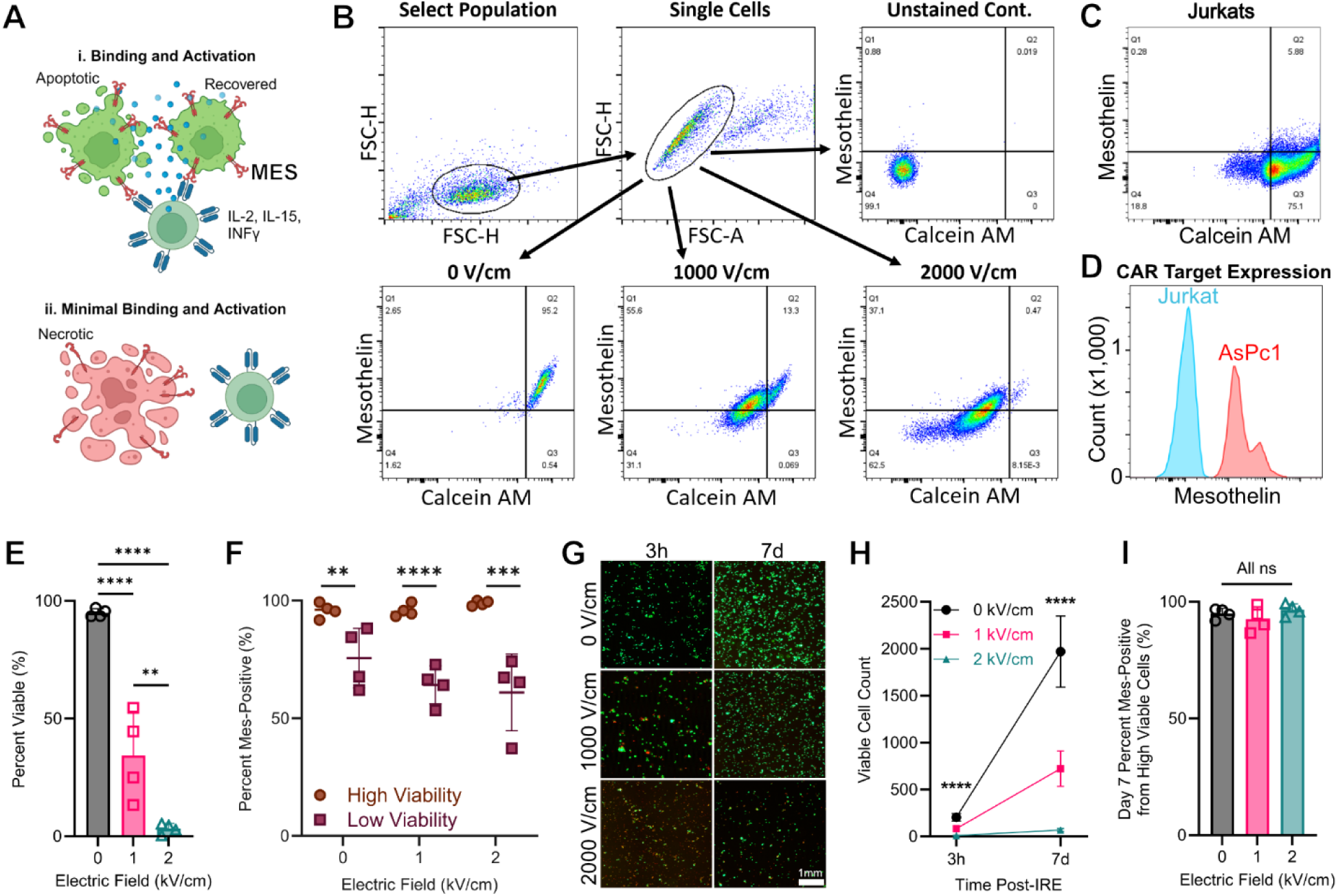
CAR target binding analysis following irreversible electroporation. **A)** Viable and intact cells following electroporation are still available for cell membrane mesothelin (MSLN) binding, while necrotic cells experience a decrease in binding. **B)** Flow cytometry gating to isolate single cells and plots of Calcein AM versus mesothelin at 3 hours following IRE delivery using 0 (control), 1000, and 2000 V/cm. **C)** Control using mesothelin-negative Jurkats. **D)** Mesothelin expression in AsPC-1s compared to Jurkats. **E)** Cell viability 3 hours after electroporation at different electric field strengths; one-way ANOVA with Tukey’s post- test and correction; mean ± SD; n=4. **F)** Percent mesothelin expression for high viability and low viability cell populations 3 hours after electroporation; multiple two-tailed T-test; mean ± SD; n=4. **G)** Live (green) and dead (red) imaging at 3 hours and 7 days after treatment; scale bar is 1 mm. **H)** Viable cell count at different electric fields after IRE and following recovery; one-way ANOVA with Tukey’s post-test and correction within each timepoint; mean ± SD; n=4. **I)** Mesothelin binding for recovered cells at day 7; one- way ANOVA with Tukey’s post-test and correction within each timepoint; mean ± SD; n=4. ***\**** *p <* 0.05*, ** p <* 0.01, *** *p<0.001*, **** *p<*0.0001.

We next treated AsPC-1s cell suspensions with conventional IRE pulses at different electric fields to induce significantly different brackets of percent viability (Figure 2E), with no death in the negative control (94.99 ± 1.74 [93.53 – 97.10] %), moderate cell death at 1 kV/cm (34.40 ± 18.65 [13.37 – 54.59] %), and almost complete cell death at 2 kV/cm (3.30 ± 2.19 [0.47 – 5.40] %). The percent high-mesothelin expression was then determined within the high-viability and low-viability populations (Figure 2F). For each treatment magnitude group, the high-viability population had a significantly higher mesothelin expression. However, when examining either the high-viability or low-viability populations, no significant differences in mesothelin expression were observed between the treatments. Subsets of the treated cells were then plated to allow for the viable cells to rebound over 7 days (Figure 2G). The viability qualitatively matched the trends observed for the percent viability of cells observed with flow cytometry at 3 hours post-treatment. For all three groups, a portion of the treated cells rebounded (Figure 2H). Analyzing the viable cells indicated high mesothelin expression in the high-viability population that rebounded following treatment (Figure 2I), with no significant difference between treatment groups within the high-viability populations at 3 hours or 7 days after IRE. These data suggest that residual cancer cells surviving IRE treatment present proteins that chimeric antigen receptors can recognize.

### An in vitro assay for monitoring treatment response and tumor recurrence following irreversible electroporation

While the protein targets for CAR binding are still functional in viable cells following electroporation, it is necessary to validate that the target protein is recognizable by CAR T-cells for subsequent removal. Here, we employed a tumor spheroid viability assay as previously described for high- throughput CAR T-cell analysis^89^, with modification to include irreversible electroporation treatments within the pipeline (Figure 3A). *In vitro* tumor spheroids provide a bridge between computational modeling and *in vivo* validation to determine treatment efficiency and explore biological mechanisms in a reproducible and high-throughput system^90,91^. 3D cultures have emerged to achieve higher biomimicry of cell–cell and cell-ECM interactions^92,93^, gene expression, cellular heterogeneity, and microarchitecture of the native tissue^94–96^. Consequently, 3D models improve the predictability of therapeutic toxicity and sensitivity, with significantly higher drug resistance observed in 3D models compared to 2D models^97^. Given their unique advantages, developing in vitro 3D models has been of growing interest within adoptive cell therapy development for assessing target binding affinity, comparing treatment efficacies, and determining patient-specific treatment responses before clinical delivery^98^.

**Figure 3.**
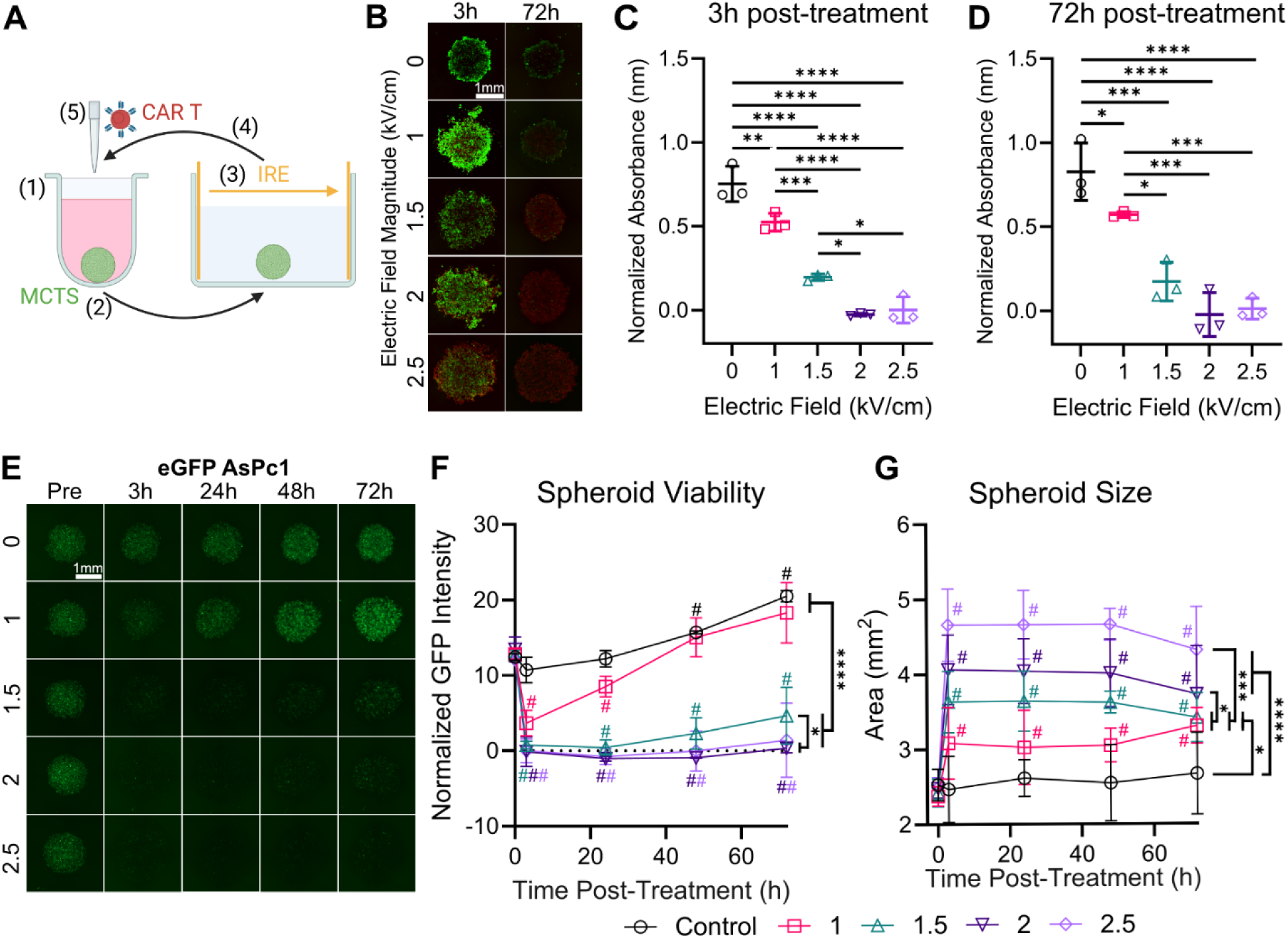
*In vitro* assay for longitudinal combinatorial treatment evaluation. **A)** *In vitro* tumor spheroid (TS) assay to assess the treatment response to electroporation and CAR T-cell therapy. (1) TSs were formed within a low-adherent U-bottom 96-well, (2) then moved to a 4-well rectangular plate with low- conductivity buffer to (3) deliver electroporation via parallel plate electrodes. (4) The TSs were immediately moved back into the original U-bottom well, where (5) adjuvant CAR T-cells therapy or sham was delivered. **B)** Live (green) and dead (red) imaging of AsPC-1 TSs at 3 hours and 72 hours after treatment for different electric field magnitudes; scale bar is 1 mm. Normalized absorbance for the XTT assay at **C)** 3 hours and **D)** 72 hours post-treatment at different electric fields; one-way ANOVA with Tukey’s post-test and correction; mean ± SD; n=3. **E)** Green fluorescent intensity of FLuc-eGFP^+^ AsPC-1 TSs over time across different electric fields. **F)** Normalized GFP intensity and **G)** TS area over time; One-way ANOVAs with Tukey’s post hoc between groups on the last timepoints (*p < 0.05, *** p < 0.001, **** p<0.0001); Multiple One-Sample Wilcoxon signed-ranked tests between that timepoint and zero (# p < 0.05); n = 3.

Multicellular tumor spheroids (TS) were fabricated using the MSLN^+^ AsPC-1s, and an initial study was performed to evaluate the treatment response to IRE at different electric field strengths. Similar to the suspension and flow cytometry data, the live/dead staining revealed a stark decrease in viability at 3 hours post-treatment that was completely resolved at 72 hours post-treatment (Figure 3B). To validate that the observed trends in viability, we utilized a quantitative XTT (sodium 3’-[1-(phenylaminocarbonyl)-3,4-tetrazolium]-bis(4-methoxy-6-nitro) benzene sulfonic acid hydrate) assay. The cells reduce the tetrazolium salt to a formazan product, which is directly proportional to the number of viable and metabolically active cells. There was a significant decrease in metabolic activity for all treatment groups (1 – 2.5 V/cm) when compared to the control, at both 3- and 72-hours post-treatment (Figure 3C-D). The metabolic activity of TS treated with 1 V/cm and 1.5 V/cm was significantly different than every other group at 3 hours, but the metabolic activity for the 1.5 V/cm treatment significantly decreased by 72 hours, with no significant differences to the 2.0 V/cm and 2.5 V/cm groups. These data support that cell death and recovery following IRE are dynamic, providing rationale for the need to track treatment response at multiple time points. However, the XTT assay has limitations for tracking spheroid viability with cellular therapies, as the measured metabolic activity will be influenced by the activation and proliferation of CAR T-cells, especially with concurrent cell death following IRE.

To enable longitudinal and isolated tracking of TS response with CAR T-cell treatments, we transfected the MSLN^+^ AsPC-1s with a firefly luciferase–enhanced Green Fluorescent Protein (FLuc-eGFP) lentivirus (Figure 3E). We observed a stark decrease in eGFP fluorescent intensity as soon as 3 hours following IRE for all treatment groups compared to the control, and we continued to measure the average GFP intensity within the TS to quantify viability for up to 72 hours (Figure 3F). The GFP intensity at 3 hours dropped significantly from baseline for all treatment groups. However, for 1 and 1.5 kV/cm, the GFP intensity then increased over the following 69 hours, with significantly higher GFP intensities at 72 hours compared to 3 hours, indicating incomplete treatment and proliferation of surviving cells within the tumor spheroid. The 2 and 2.5 kV/cm groups did not significantly change from their initial drop at 3 hours. This data validated the viability data observed from the live/dead imaging and metabolic assays, providing a novel method for tracking individual tumor spheroid viability and proliferation for longitudinal electroporation studies.

Following treatment, the size of the TS significantly increased for every IRE group compared to their pre-treatment measurements (Figure 3H). The spheroid size did not change significantly for the control across all time points and remained unchanged for all IRE groups after the initial significant increase in size measured at 3 hours. Further, the change in spheroid size increased significantly with higher applied electric fields. Since the metabolic activity decreased with increased electric field and the number of cells within the spheroid was equal across all replicates, the increase in TS size cannot be attributed to proliferation. These data suggest that IRE modulates the TS by decreasing cellular density, potentially through the dysregulation of cell-cell adhesion in low-viability cells, as supported by the decrease in cell surface protein binding observed for the MSLN receptor.

### In vitro spheroid study supports that CAR T-cell therapy can eliminate residual cancer following irreversible electroporation

Utilizing the TS assay, we observed that the cellular organization was disrupted following IRE, and proliferation was preserved in portions of the TS for applied electric fields below 1500 V/cm. To investigate whether CAR T-cell therapy can eliminate residual cancer and whether the lower density facilitates CAR T-cell penetration into the TS, we utilized our FLuc-eGFP^+^ AsPC-1 TS assay again, with the addition of adjuvant CAR T-cells (Figure 4A-C). CAR T-cells were fabricated using primary human T-cells as previously described^99–102^ to express a 2^nd^ generation anti(α)- human(hu)MSLN-m28-mCD3z CAR gene against huMSLN^102–104^. The size of the spheroids treated with only CAR T-cells decreased significantly over time, in contrast to the controls, which remained steady (Figure 4D). The size of the spheroid again increased significantly at 3 hours following 1500 V/cm IRE due to dysregulation of the TS. However, with the subsequent addition of CAR T-cells, the size then decreased significantly over time, suggesting a potential combinatorial effect on the cells within the TS. The improved cytotoxicity from the combinatorial therapy was further supported by the normalized GFP intensity within the spheroid (Figure 4E). The GFP intensity within the combinatorial treatment dropped by 3 hours and remained low, with a significantly lower GFP intensity than the other groups, including the IRE-only group that showed evidence of partial recurrence by 72 hours. Although the size of the TS decreased over time within the CAR T-cell group, the fluorescent intensity was not significantly different from the control until 72 hours, when the GFP intensity decreased significantly. This suggests that the CAR T-cells are binding and inducing cytotoxic effects on the outer layers of the TS without deeper penetration until 72 hours. To investigate CAR T-cell localization within or around the TS, we stained the CAR T-cells with CellTracker^TM^ deep-red. Using the eGFP images, we defined a region of interest outlining the TS border and then overlaid it on the cell tracker deep-red images to quantify the normalized deep-red intensity within the TS (Figure 4E). The deep-red intensity was initially significantly higher within the TS following IRE. Measuring over 72h, the intensity remained unchanged within the combinatorial group but significantly increased within the CAR T-cell only group (Figure 4F). The increase in infiltration by 72 hours within the CAR T-cell only group corresponds with the decrease in spheroid size and decrease in viability at 72 hours, verifying the successful generation of anti-tumor CAR T-cells and validating cytotoxicity against cancer cells expressing bindable CAR targets. The initially higher infiltration within the combinatorial group suggests that the disrupted TS allows for the penetration of CAR T-cells; however, due to the significantly lower viability and subsequent lower presence of bindable CAR targets, the CAR T-cells do not have pressure to invade. Further, the data indicated that there is an increase in efficacy of combinatorial IRE and CAR T-cell therapy, preventing the few viable cells from repopulating the TS, and that many of the CAR T-cells within the combinatorial group did not need to invade for the additional cytotoxic effects.

**Figure 4.**
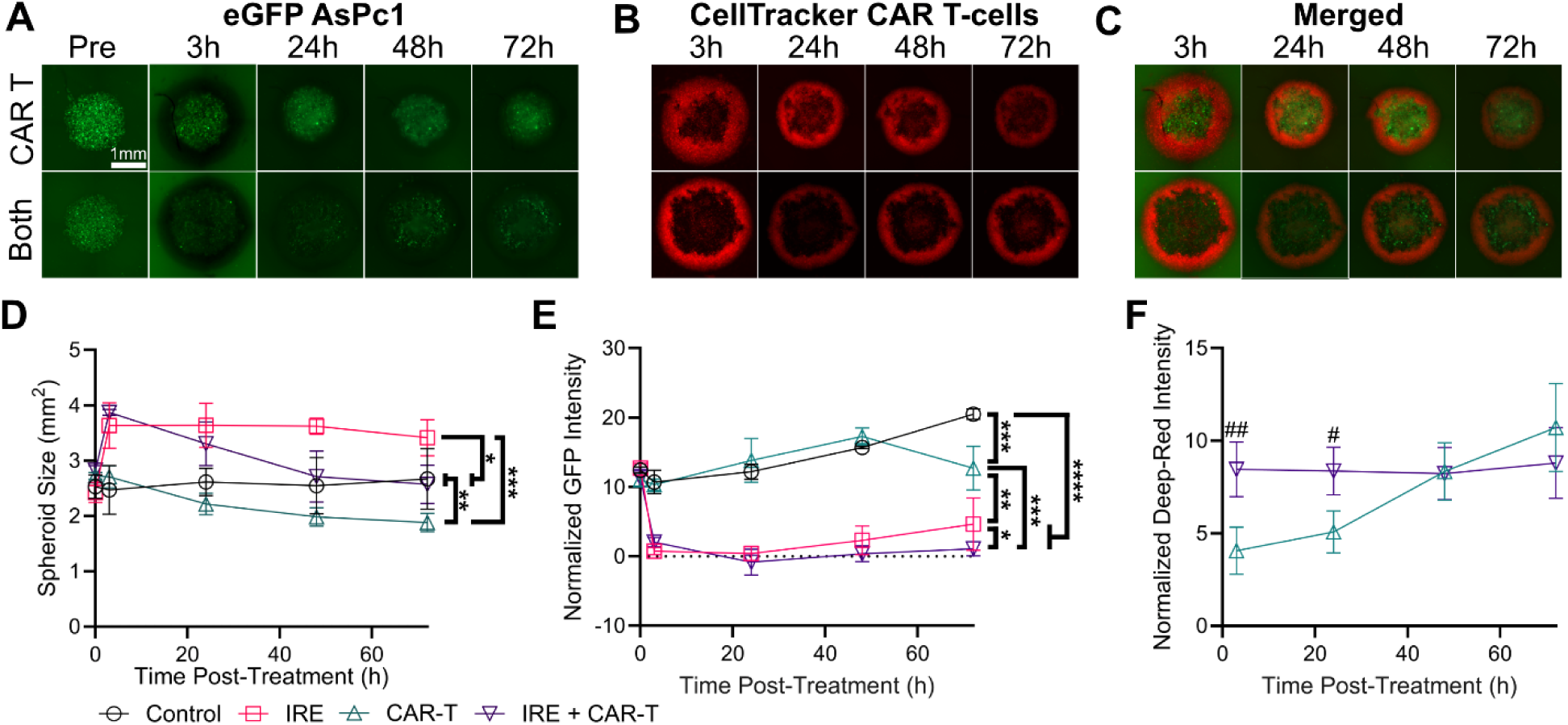
*In vitro* evaluation of anti-tumor efficacy and infiltration following IRE and CAR T-cell therapy. **A)** Green fluorescent intensity of FLuc-eGFP^+^ AsPC-1 TS, **B)** deep-red intensity of cell tracker-stained CAR T-cells, and **C)** merged image over time for the CAR T-cell only and combinatorial treatment (both). **D)** Measured TS area and **E)** normalized eGFP intensity over time; One-way ANOVAs with Tukey’s post hoc between groups on the last time points (*p < 0.05, ** p< 0.01, *** p < 0.001, **** p<0.0001); n ≥ 3. **F)** Comparison of deep-red intensity within the tumor spheroid over time; Two-tailed T-tests between groups at each timepoint (# p < 0.05, ## p < 0.01), n ≥ 3.

### Combinatorial irreversible electroporation and CAR T-cell therapy have enhanced efficacy in a pre-clinical human pancreatic cancer mouse model

While *in vitro* studies are essential for evaluating novel CAR designs, the limitations for CAR T-cells are well-documented *in vivo* and cannot be fully recapitulated *in vitro*^73,86,98^. Immunotherapies, including CAR T-cell therapies, are also required to be evaluated *in vivo* for efficacy and are shown to be mouse strain dependent, highlighting the need for appropriate strain selection for specific studies^105–107^. Severely immunodeficient NSG mice are the most common model in adoptive cell therapy research, as they can engraft human cancers, enabling the development of patient-specific therapies^105^. To validate the proof-of-concept combination of IRE and CAR T-cell therapy demonstrated using the *in vitro* TS assay, we inoculated NSG mice with 1x10^6^ of the non- transfected MSLN^+^ AsPC-1 cells (Figure 5A). The AsPC-1 tumors grew considerably slower than the Pan02s, reaching 38.59 ± 10.68 [22.5 – 54.675] mm^3^ in volume at 23 days. The tumors were treated with IRE as detailed above (Figure 5B), with immediate local injection of 5x10^6^ αhuMSLN CAR T-cells. To visualize CAR T-cell localization and verify successful transfection, the CAR gene is concomitantly expressed with firefly luciferase. IVIS imaging, both before and after treatment, verified the successful delivery of CAR T-cells into the tumor (Figure 5C). Body weight measurements did not significantly change after tumor inoculation until euthanasia (Figure 5D), and wellness checks by the researchers and the department of animal research staff did not indicate cytotoxicity associated with treatment. These data and observations suggest that the combinatorial treatment was well tolerated. Two of three tumors (66.6%) within the combinatorial treatment group were eradicated at 7 days post-treatment, whereas the control, standalone IRE, and standalone CAR T-cell groups still had measurable tumors. The tumor volumes 7 days post-treatment were not significantly different between the combinatorial treatment, standalone CAR T-cell treatment, and standalone IRE treatment, but all three groups were significantly different than the control (Figure 5E). When comparing the percent change in tumor volume, the combinatorial treatment had a significantly higher reduction than the other groups (Figure 5F). These data validate the *in vitro* results, indicating an increased therapeutic efficacy for the combination of IRE and CAR T-cell therapy.

**Figure 5.**
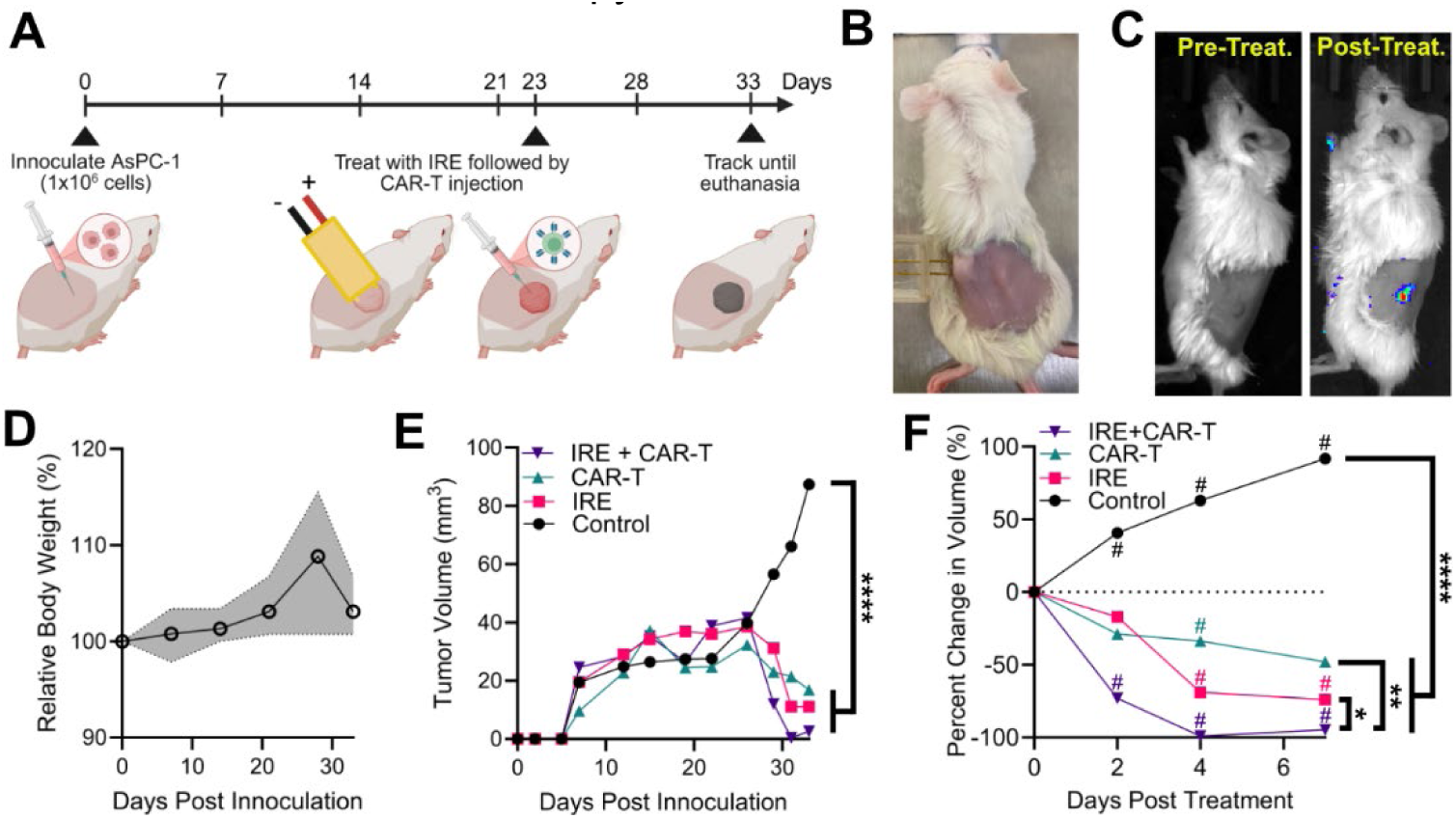
Combinatorial Treatment. **A)** Schematic of the treatment timeline: NSG mice were inoculated with MSLN^+^ AsPC-1s on day 0. Mice received IRE followed by CAR-T injection on day 23 and were tracked for 7 days until day 33. **B)** Image of IRE delivery through 2-needle electrodes. **C)** Pre- and post-treatment IVIS-imaging reveals FLuc^+^ αMLSN CAR T-cells at the tumor site. **D)** Average and range of rodent weights; Multiple paired T-tests between the data compared to the initial day zero weight; n=8 **E)** The average measured tumor volume from inoculation and **F)** average percent change in tumor volume after treatment on day 23 post inoculation; One-way ANOVAs with Tukey’s post hoc between groups on the last timepoints (***\*****p < 0.05, ** p < 0.01, **** p<0.0001*); Multiple One-Sample Wilcoxon signed-ranked tests between that timepoint and zero (# *p < 0.05*); n = 3 for control and IRE + CAR-T, n=2 for standalone IRE and CAR-T.

## Discussion

Here, we demonstrate the scientific and clinical feasibility of a first proof-of-concept combination of irreversible electroporation with engineered cells and support the use of pulsed field ablation techniques with an adoptive cell therapy, specifically CAR T-cell therapy. Our findings demonstrate that IRE does not compromise the structural integrity of membrane-bound tumor antigens in cells that survive treatment, thereby preserving the binding potential of chimeric antigen receptors. This mechanistic insight is critical, as other ablation techniques use thermal energy, which can indiscriminately denature proteins, potentially limiting compatibility with targeted immunotherapies. In contrast, IRE uniquely retains epitope integrity in viable cells within the treatment margin, enabling CAR T-cells to recognize and eliminate residual disease. This highlights the potential for IRE to serve not only as a cytoreductive tool for otherwise unresectable tumors but also as a unique facilitator of antigen-targeted immune engagement in solid tumors commonly treated by thermal ablation. Nevertheless, mesothelin is a cell surface glycoprotein with a heavy and light chain, which may not denature as easily from high electric fields as other CAR targets that are complexes comprised of many constitutive parts (e.g., HER2, IL13Ra2). Thus, similar evaluations used to investigate mesothelin binding through flow cytometry should be performed to verify recognition of other CAR targets following electroporation.

We contend that IRE and CAR T-cell therapy may also synergize their treatment pipelines. Many patients receive chemotherapy before irreversible electroporation^34,39,65^, so T-cells can be removed to manufacture CAR T-cells *ex vivo* for reinfusion at treatment. This combination can then overcome the aspects of a weakened anti-tumor immune response due to lymphodepletion in these patient populations^77^. Further, cytopenias and neurotoxicity are also directly correlated to CAR T-cell concentration^73^, which can limit the efficacy of the therapy, as high concentrations are typically required for solid tumors. A previous study found that 5x10^6^ MSLN CAR-Ts alone could not clear the primary pancreatic tumor in NSG mice but significantly prevented the initiation and progression of lung metastases^108^, informing the concentration used for this proof-of-concept combination of IRE with CAR T-cell therapy. We found similar results for the standalone CAR T-cell group, with the tumor size only significantly decreasing from the pre-treatment size at day 4 post-treatment. The combination of IRE and CAR T-cell therapy resulted in a reduction in tumor size, leading to the removal of measured tumors compared to either standalone IRE or CAR T-cell therapy. The *in vitro* assay further supported the increased efficacy and cytotoxicity of the combinatorial treatment, but it also revealed that CAR T-cell engagement with the TS was lower in the IRE-treated TS, potentially due to the reduction in MSLN binding in low-viability cells. These data suggest that IRE can act as a cytoreductive technique for reducing the dose of adoptive cell therapies required for therapeutic efficacy. Future work should investigate the therapeutic CAR T-cell dose curve with and without concomitant IRE to verify whether equivalent efficacies can be achieved with fewer CAR T-cells, potentially alleviating severe side effects attributed to targeted therapies.

Antigen escape is a potential mechanism by which tumor cells evade CAR T-cell therapy^72,73,86^, which IRE may indirectly address. While there was a significant reduction in tumor size for the combinatorial treatment, a few cells could evade both therapies and the adaptive immune response to repopulate the tumor. Tandem CAR T-cells aim to overcome this by targeting multiple antigens simultaneously, decreasing the chance that cells can downregulate both targets. Dual and triple CAR T-cells have been demonstrated to increase the therapeutic efficacy but have the chance to significantly increase on-target off-tumor cytotoxicity^109^. Reducing the necessary CAR T-cell concentration with cytoreductive IRE may allow for safe use of tandem CAR T-cells. Patient-derived xenografts with heterogeneous antigen expression or mixtures of antigen-positive and antigen-negative cells should be utilized to evaluate the potential for complete tumor eradication with combinatorial treatment.

The application of either IRE or CAR T-cells as a monotherapy has proven successful in enabling an endogenous anti-tumor response through the induction of pro-inflammatory cytokines, facilitating local immune cell recruitment. Additionally, IRE has been shown to activate the systemic anti-tumor response, resulting in a reduction of distant metastasis in rodents^44^, similar to the “abscopal effect” that is rarely observed in patients exposed to radiotherapy. With the sequential delivery of IRE and CAR T-cells, the combinatorial treatment offers the potential to more consistently generate a robust local and systemic immune response that can persist beyond the initial treatment by priming the adaptive immune response. An adaptive immune response may then recognize the cancer that has downregulated the CAR target. While this work demonstrated proof-of-concept for treating human disease, future work should utilize a synergistic tumor and allogenic CAR T-cell combination within an immunocompetent model to investigate the use of IRE for facilitating the infiltration and persistence of adoptive cell therapies in an immunogenically active TME (Figure 6A). We expect that CAR-T infiltration and persistence will be significantly improved following IRE, with increased recruitment of the endogenous immune system. This may then “wake up” a potentially dysregulated immune response to respond to local and metastatic disease (Figure 6B).

**Figure 6.**
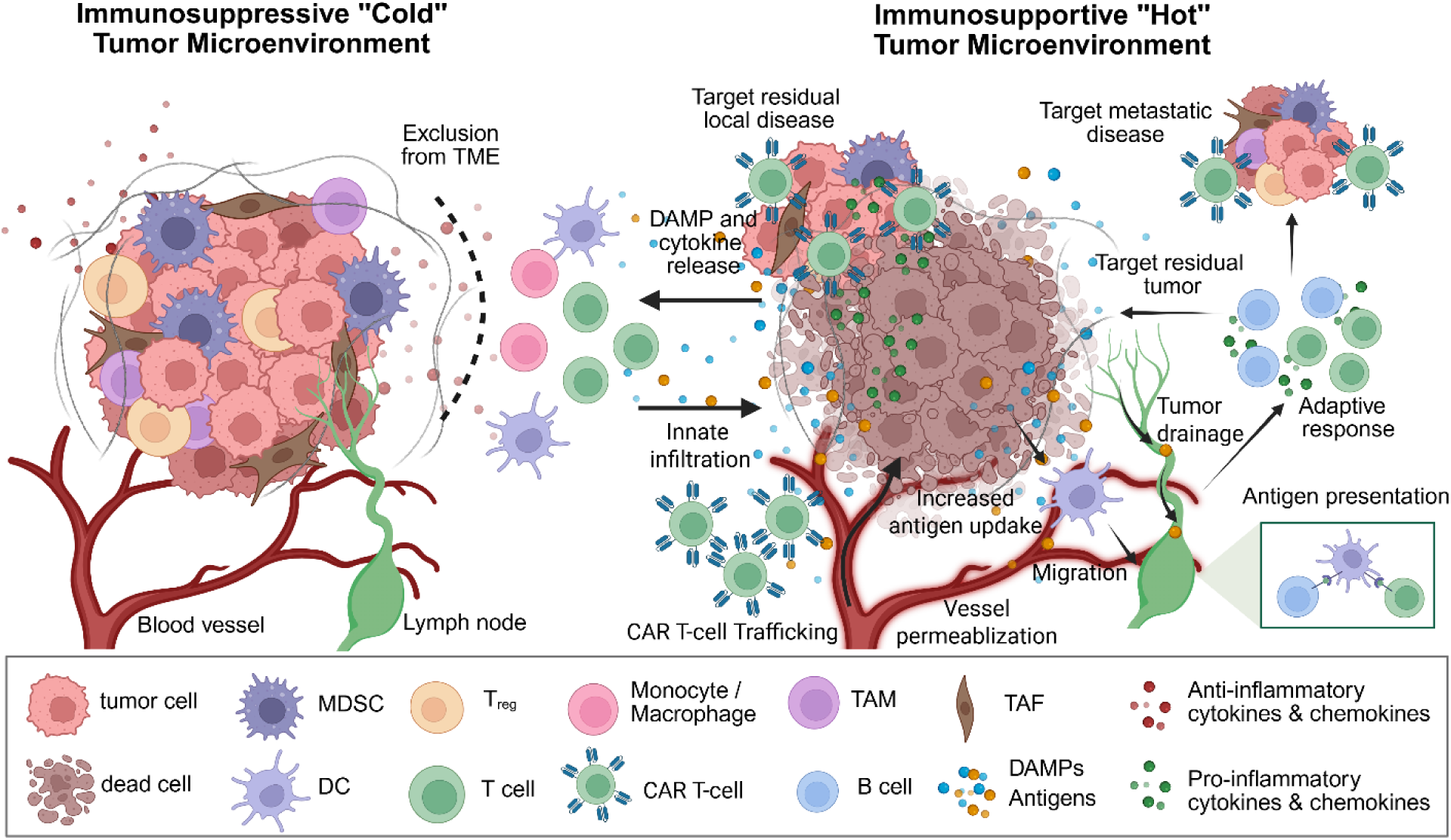
Summary and Future Directions. **A)** IRE indiscriminately ablates cells within the dense and immunosuppressive tumors, clearing the regulatory T cells (T-regs), tumor-associated macrophages (TAMs), myeloid-derived suppressor cells (MDSCs), and the bulk tumor. **B)** Post-ablation, systemic CAR T-cell therapy can then effectively persist within the tumor microenvironment to clear the residual tumor. Adapted from Jacobs *et al.* 2025^14^ (Creative Commons Attribution 4.0 International License).

The AsPC-1 tumors grew considerably slower than the Pan02s for the same number of cells inoculated per tumor, with the AsPC-1 control tumors not reaching 100 mm^3^ by the endpoint. Future experiments should allow the AsPC-1s tumors to grow longer or conduct a similar recurrence study using luminescent reporter AsPC-1s. While we found significant differences in treatment responses for the combinatorial treatment, the limited sample size reduced the statistical power. Larger longitudinal studies are required to validate the findings presented and to evaluate the TME itself, thereby determining the reasons for the improved treatment response.

## Conclusions

This study presents the first demonstration of combining irreversible electroporation (IRE) with chimeric antigen receptor (CAR) T-cell therapy for the treatment of solid tumors. In immunodeficient mouse models bearing human pancreatic tumors, we confirm that IRE alone may leave behind viable residual disease, underscoring the need for immune-mediated clearance. We demonstrate that IRE preserves the structural integrity of cell surface antigens on viable tumor cells, thereby enabling effective CAR T-cell binding and cytotoxicity following ablation. Our *in vitro* and *in vivo* findings support the feasibility and potential synergy of this dual-modality approach; IRE disrupts the immunosuppressive tumor microenvironment while reducing tumor burden, thereby enhancing the accessibility and efficacy of adoptive cell therapies. This combinatorial strategy addresses critical barriers facing CAR T-cell therapies in solid tumors, establishing a foundation for future investigations into dosing, timing, and immune integration in clinically relevant models. Together, these results demonstrate a novel therapeutic paradigm for pancreatic cancer that integrates broad focal ablation and precise patient-specific immunotherapy.

## Methods

### Cell culture

Pan02 mouse pancreatic cancer cells (Cytion, 300501), AsPC-1 human pancreatic cancer (CRL-1682), and Jurkat immortalized human T-lymphocytes (ATCC, TIB-152) were cultured in RPMI-1640 medium (ThermoFisher, 11875093) supplemented with 10% (v/v) fetal bovine serum (Fisher Scientific, FB12999102) and 1% (v/v) 10,000 U/ml penicillin-streptomycin (Gibco, 16140122). The cells were maintained at 37°C and 5% CO2 in a regulated, humidified incubator (Fisher Scientific, 13-998-086). Pan02s and AsPC-1s were passaged between 70 and 90% confluency, and Jurkats were maintained in cell suspension between 1x10^6^ and 2x10^6^ cells/ml. All cells were utilized between passages 10 and 20.

### Pan02 transfection with reporter firefly luciferase (FLuc) – enhanced Green Fluorescent Protein (eGFP)

Pan02s were plated at 1x10^4^ per well in a 96-well plate and incubated overnight. A 2-fold titration of a FLuc–eGFP lentivirus (BPS Biosciences, #79980), driven under a single cassette by the CMV promoter for co-expression, was performed from a max MOI of 100, with the lentivirus diluted in supplemented cell culture media. The cells were incubated for 8 hours, then the media-lentivirus mixture was replaced with fresh supplemented cell culture media. The cells were incubated for 3-5 days until confluent, with transfection confirmed through green fluorescent from eGFP using a fluorescent microscope (Leica, LMI8). The cells were then fully passaged into a 6-well plate for expansion. Once confluent, the cells were released and suspended in 10 ml of supplemented media for cell sorting using a 9-color FACS Melody^TM^ cell sorter (BD Biosciences). Following gating for the cells and isolation of single cells, the cells were gated for high eGFP expression, and the population was collected. The cells were then resuspended for cell culture. Stocks of FLuc-eGFP Pan02s were frozen in liquid nitrogen.

### NOD/SCID/IL2gc-KO (NSG) mice

NSG mice were obtained through breeding at the Georgia Tech Department of Animal Research (DAR), with ages ranging from 16 to 24 weeks at the start of experiments. All rodent experiments were conducted at the DAR under IACUC approval (protocol no. A100702) at Georgia Tech and in accordance with the NIH Guide for the Care and Use of Laboratory Animals. Mice were maintained under sterile housing with food, water, and cage changes occurring under fumigation.

### Pan02-NSG local tumor recurrence experiments

FLuc-eGFP Pan02s were washed 2x with sterile HBSS (ThermoFisher, 14025092), resuspended at 20x10^6^ cells/ml, then mixed with an equal volume of sterile Matrigel (Corning™ Matrigel™ Matrix, CB-40234C). At the GT DAR, NSG were subcutaneously inoculated with 100 μl of the cell suspension after the site was shaved and sterilized with an isopropyl wipe. Injection sites were measured three times a week to monitor tumor size. Mice were treated once the tumors reached >100 mm^3^, where 𝑣𝑣𝑣𝑣𝑣𝑣𝑣𝑣𝑣𝑣𝑣𝑣 = 0.5 · 𝑣𝑣𝑣𝑣𝑙𝑙𝑙𝑙𝑙𝑙ℎ · 𝑤𝑤𝑤𝑤𝑤𝑤𝑙𝑙ℎ^2^. Prior to irreversible electroporation (IRE) pulse delivery, the mice were initially imaged using the *in vivo* imaging system (IVIS) at the GT DAR. Briefly, 10 μl of a 15 mg/ml D-luciferin (GoldBio, LUCK-1G) in sterile HBSS was injected intraperitoneally (*i.p.*), with serial imaging performed every 2 minutes for 45 minutes to determine the optimal imaging time window. Following imaging, irreversible electroporation was delivered through custom 2-needle electrodes, as previously described^78,79^, using a custom pulsed electric field generator (Vitave Inc., OmniPorator, Czech Republic). The conventional ninety bursts of 90-μs pulses were applied at a rate of 1 Hz. The applied voltage was adjusted to deliver 2500 V per cm of center-to-center electrode spacing, with the spacing set to be ∼1 mm less than the max tumor diameter to facilitate insertion into the tumor. The applied voltages and currents were recorded using a WaveSurfer 5 GHz oscilloscope (Teledyne LeCroy, 4024HD) equipped with a 1000× attenuated high-voltage probe (Siglent, DPB5700) and a 10× attenuated current probe (Pearson Electronics, 3972). Following treatment, the needle insertion sites were inspected for potential bleeding, and the animals were recovered. Mice were then again imaged using IVIS on day 3, day 7, and day 17. *In vivo* imaging luminescent data were collected and analyzed with living image 4.4.5 (PerkinElmer).

### Mesothelin binding and viability following electroporation

1x10^6^ AsPC-1s were suspended in osmotically-balanced low conductivity buffer^110^, and 100 μl was pipetted into a sterile 4-mm cuvette (BTX, #45-0142). Pulsed electric fields were delivered using the custom generator and pulsing scheme described above at 0 V, 400 V, and 800 V to generate electric field magnitudes of 0 V/cm, 1000 V/cm, and 2000 V/cm, respectively. The treated cells were then either transferred to a 24-well plate with fresh cell culture media or to a 1.5-ml tube with 400 μl of cell culture media for flow cytometry. For the plated cells, the cell culture media was replaced on day 3, and at day 7, they were released to resuspend in media in a 1.5-ml tube for flow cytometry. The processing and flow cytometry steps were identical for both the 3-hour and 7-day time points, with the timing for the 3-hour group adjusted so that flow cytometry occurred at 3 hours following treatment. Cell suspensions were centrifuged at 500 x g, with the liquid aspirated, and resuspended in 500 μl in flow cytometry buffer (ThermoFisher, 00-4222-26). The cells were again centrifuged and resuspended with 1 μl (10 μl / 10^6^ cells) Human Mesothelin Alexa Fluor^®^ 488-conjugated Antibody (RNDSystems, FAB32652G) and 0.5 μl Invitrogen™ CellTrace™ Calcein Red-Orange, AM (Fisher Scientific, 50-113-7411) in 500 μl of flow cytometry buffer. Tubes were placed on a tube rocker on low within a dark 4°C refrigerator for 45 minutes. The cells were then washed twice with flow cytometry buffer to remove excessive stain, resuspended in 500 μl of flow cytometry buffer, and then analyzed using a Cytek^®^ Aurora (Cytek Aurora). Flow cytometry data were collected with BD FACSDiva v8 (BD Biosciences, San Jose) and analyzed using FlowJo X v11 (FlowJo, Ashland). Plated cells not used for flow cytometry were stained for viability at 3 hours and 7 days following treatment. Cell culture media was replaced with a live/dead stain consisting of 0.5 μl Invitrogen™ CellTrace™ Calcein Green, AM (ThermoFisher, C3100MP) and 2.5 μl propidium iodide (ThermoFisher, P3566) in 250 μl of phosphate buffered saline (ThermoFisher, 10010023), then imaged using a Leica DMI8 fluorescent microscope. Cells were counted within the Leica LasX software within the region of interest corresponding to the image gathered using the 10x objective.

### Chimeric Antigen Receptor (CAR) T-cell fabrication

Human T cells were isolated from peripheral blood mononuclear cells (PBMCs) of a healthy donor and activated using αCD3/CD28 Dynabeads (ThermoFisher, # 11132D) and 200 U/mL hIL-2 for 24 hours. During the activation, T cells were transduced with hMSLN- (M5)-h28Z-Luc-CAR encoding γ-retrovirus^111^ to make T cells targeting human MSLN by centrifugation at 1,800 RPM for 2 hours at room temperature. T cells were then expanded in the presence of 200 U/mL hIL- 2 for an additional 7–8 days before use.

### Tumor spheroid experiments

To create tumor spheroids (TS), either AsPC-1 or FLuc^+^eGFP^+^ AsPC-1s cells were passaged and resuspended in supplemented cell culture media at 5x10^5^ cells/ml, with 100 μl pipetted into a 96-well ultra-low adherent (ULA) plate. The plate was centrifuged at 150 x g for 5 minutes, and the cells were monitored for 2 days to verify TS formation. To apply IRE, the spheroids were removed and placed in a rectangular 4-well plate (Ibidi) containing 400 μl of low-conductivity buffer. Custom flat plate electrodes with an 8.5 mm spacing were used to deliver uniform pulsed electric fields, with the voltages and currents monitored as described above. The TSs were then placed back into the ULA plate with fresh 200 μl of supplemented cell culture media. CAR T-cells were stained for 30 minutes with 1μM CellTracker^TM^ Deep-Red (ThermoFisher, C34565). For groups receiving CAR T-cell treatments, CAR T-cells were added at a 5:1 ratio to the pre-treatment AsPC-1s cell count (i.e., 25x10^5^ cells). The viability of non-transfected AsPC-1s was imaged using the live/dead stain detailed above. The XTT assay was performed according to the manufacturer’s instructions; 50μl of the XTT-activated assay reagent was added to each well and incubated for 18 hours. The supernatant within each well was then transferred to a 96-well plate, and absorbance was recorded at 475 and 660 nm (Biotech Synergy HT). The measured absorbance at 475 nm was normalized using the readings at 660 nm. Size and fluorescent intensity of the FLuc^+^eGFP^+^ AsPC-1s TSs were quantified within ImageJ (NIH). Each image was separated into the individual channel components (i.e., bright field, green, and deep-red). A region of interest, informed by bright field imaging and green fluorescence, was drawn around the border of the TSs. The intensities of both the green and deep-red channels were measured within the TS and outside to obtain background intensity, with the color intensities normalized by subtracting the background intensities.

### AsPC-1 NSG mouse experiments

1x10^6^ AsPC-1s were prepared and subcutaneously inoculated in NSG mice as described above for the Pan02-NSG experiment, with the injection sites measured three times a week to monitor tumor size. Mice were randomly grouped and treated at day 23 with either (1) sham electrode insertion with local saline injection, (2) IRE with local saline injection, (3) sham electrode insertion with local CAR T-cell injection, or (4) combinatorial treatment using IRE with local injection of CAR T-cells within 1 minute following treatment. IRE delivery was performed as detailed above. The local injections were performed slowly to prevent backflow, and for CAR T-cell delivery, 5x10^6^ cells were injected in 100 μl sterile saline. The mice were imaged before and after treatment using IVIS. Following treatment, the needle insertion sites were inspected for potential bleeding, and the animals were recovered.

### Statistical Analyses

Statistics within the main text are presented as mean ± SD [minimum – maximum]. Statistical analyses were performed using GraphPad Prism 10.5.0 (GraphPad Software, Boston), with the specific tests used outlined within the text. Post-experimental power analyses were performed using G*Power 3.1 (Heinrich Heine Universität, Düsseldorf).

## Funding and Acknowledgments

This research was partly funded by the NIH R01 CA240476 and NIH R01 CA274439. We would like to acknowledge Jessica Weems at the Georgia Tech Department of Animal Research breeding program for their donation and help with mouse housing, Erich Williams at the Georgia Tech Cellular Analysis and Cytometry Core for discussions and help with flow cytometry data generation and analysis, and Michal Cifra at the Institute of Photonics and Electronics of the Czech Academy of Sciences for early discussions on protein analysis experiment design.

## Data Availability Statements

The data supporting the findings of this study are available within the article. Further requests can be made to the corresponding author.

## Disclosures

E.J.J, J.P.A., and R.V.D. have submitted patents related to electroporation and adoptive cell therapies through Georgia Tech.

## Notes

### Competing Interest Statement

The authors have declared no competing interest.

